# SLEEP CHANGES WHAT MEMORIES BECOME: DELAYED REACTIVATION REVEALS LATENT EFFECTS OF POST-LEARNING SLEEP

**DOI:** 10.64898/2026.06.23.733982

**Authors:** Malen D. Moyano, Micaela S. Lombardi, Aylin A. Vazquez Chenlo, Luis I. Brusco, Cecilia Forcato

## Abstract

Sleep is thought to promote memory consolidation through the offline reactivation and reorganization of newly acquired information. However, most studies assess memory shortly after sleep, leaving unresolved whether an initial post-learning sleep episode produces enduring modifications that influence how memories respond to later reactivation. Importantly, the absence of behavioral differences after prolonged retention intervals does not necessarily imply that sleep failed to modify the original memory. Instead, sleep-dependent changes may persist in latent forms that are not readily captured by conventional memory assessments.

Here, we investigated whether post-learning sleep produces lasting changes in declarative memories that influence their subsequent response to reactivation. In Study 1, participants learned a declarative memory task and were assigned to either a short nap, a wake condition, or an exploratory long-nap condition that included both NREM and REM sleep. Memory was assessed one week later. Despite substantial forgetting across the retention interval, no significant differences in memory performance were observed between groups. In Study 2, participants learned the same task and subsequently underwent either a short nap or wakefulness. Memory was reactivated six days after learning using an incomplete reminder previously shown to induce memory updating in human declarative memory, and memory was tested one day later. Under these conditions, participants who slept after learning showed better memory performance than wake controls. Moreover, sleep physiological measures predicted the magnitude of the post-reactivation memory benefit.

These findings suggest that post-learning sleep induces enduring modifications in declarative memories that are not readily detectable through delayed memory testing alone. Instead, these sleep-dependent changes become evident when memories are challenged through subsequent reactivation. Our results indicate that sleep-dependent consolidation influences the future expression of memory, shaping how memories respond to later reactivation experiences and providing new insight into the relationship between consolidation and reconsolidation.

## 1. Introduction

Memories are not stored in a permanent form immediately after learning. Instead, newly acquired information remains initially labile and undergoes a series of stabilization processes collectively known as memory consolidation (Dudai, 2004; Dudai et al., 2015). Consolidation encompasses both local synaptic modifications and the progressive reorganization of memory representations across distributed neural networks, ultimately promoting long-term retention and successful retrieval of stored information (Squire & Alvarez, 1995; Frankland & Bontempi, 2005; Dudai et al., 2015).

Over the last two decades, converging evidence has identified sleep as a critical physiological state for memory consolidation (Diekelmann & Born, 2010; Rasch & Born, 2013; Klinzing et al., 2019). According to the Active Systems Consolidation hypothesis, recently encoded memories are spontaneously reactivated during sleep through coordinated interactions between hippocampal and neocortical networks (Diekelmann & Born, 2010; Rasch & Born, 2013; Klinzing et al., 2019). These reactivation events are thought to occur primarily during non-rapid eye movement (NREM) sleep and are supported by the temporal coordination of cortical slow oscillations, thalamo-cortical sleep spindles, and hippocampal sharp-wave ripples (Mölle et al., 2009; Staresina et al., 2015; Latchoumane et al., 2017; Helfrich et al., 2018). Through repeated reactivation, memory representations are progressively stabilized and reorganized across distributed cortical networks, supporting their long-term retention and accessibility (Diekelmann & Born, 2010; Klinzing et al., 2019).

Behaviorally, numerous studies have shown that sleep following learning improves subsequent memory performance compared with equivalent periods of wakefulness (Diekelmann & Born, 2010; Rasch & Born, 2013). In most cases, however, memory is assessed within hours or one day after the post-learning sleep period, conditions under which sleep-related benefits are robustly detected (Diekelmann & Born, 2010; Rasch & Born, 2013). Comparatively fewer studies have examined whether an initial sleep episode exerts a lasting influence on memory across longer retention intervals, from days to months, encompassing multiple subsequent sleep-wake cycles (Gais et al., 2007). Consequently, the long-term consequences of post-learning sleep remain insufficiently understood.

This distinction is particularly relevant because memory processing continues beyond the initial post-learning period, with memories undergoing ongoing stabilization and reorganization across subsequent cycles of wakefulness and sleep (Lewis & Durrant, 2011; Dudai et al., 2015). Consequently, behavioral differences observed shortly after learning may progressively diminish over time, even when the underlying memory representations remain different. Evidence from human declarative memory studies supports this possibility. For example, memory strengthening effects induced by repeated reactivation procedures can be readily detected when memories are reactivated shortly after learning, but become undetectable when the same procedures are applied several days later (Forcato et al., 2013). Thus, the disappearance of behavioral differences over time does not necessarily imply that previous manipulations failed to modify the memory. Rather, it raises the possibility that memory modifications may persist in latent forms that are not readily captured by conventional behavioral assessments.

Beyond the progressive loss of observable behavioral differences, evidence also suggests that experiences occurring shortly after learning can alter the future response of a memory. In human declarative memory, a social stress manipulation administered shortly after learning was shown to alter the subsequent response of 7-day-old memories to repeated reactivation procedures, reinstating effects that were otherwise absent in memories of the same age (Fernández et al., 2016). These findings suggest that experiences occurring during early stages of consolidation can shape the future fate of a memory and determine its later susceptibility to modification.

A relatively small number of studies have examined the consequences of post-learning sleep beyond the first days after encoding. Most investigations assess memory shortly after sleep, when sleep-dependent benefits are most readily detected (Diekelmann & Born, 2010; Rasch & Born, 2013). As a result, comparatively little is known about whether an initial sleep episode produces enduring changes in memory representations that remain relevant after multiple subsequent sleep-wake cycles. Importantly, the absence of detectable behavioral differences after prolonged delays does not necessarily imply that sleep failed to modify the original memory. Rather, it remains possible that sleep-dependent changes persist in latent forms that are not readily captured by conventional memory assessments.

One possibility is that sleep does not merely influence the immediate strength of a memory but instead modifies the state of the memory itself (Lewis & Durrant, 2011; Dudai et al., 2015; Stickgold & Walker, 2013). Under this view, sleep may generate memories that are more stable, more resistant to interference (Ellenbogen et al., 2006), or that differ in the way they respond to future reactivation experiences, even when these differences are not reflected in overt behavioral performance. Testing this possibility requires approaches capable of probing latent properties of memory beyond simple recall accuracy (Dudai et al., 2015).

Memory reconsolidation provides a useful framework for addressing this question. Once consolidated, memories can be reactivated by reminders and temporarily return to a labile state before being restabilized through reconsolidation (Nader et al., 2000; Dudai, 2006; Nader & Hardt, 2009). Importantly, previous studies have shown that the outcome of reconsolidation depends on the characteristics of the pre-existing memory. Factors such as memory age, strength, and prior learning history can determine whether a memory undergoes labilization and how it responds to subsequent reactivation experiences (Suzuki et al., 2004; Wang et al., 2009; Forcato et al., 2011; Forcato et al., 2013). Furthermore, post-learning experiences can influence the later response of a memory to reactivation procedures (Fernández et al., 2016). Together, these findings suggest that reconsolidation may serve as a sensitive tool for revealing latent differences between memories that appear behaviorally equivalent, thereby providing access to properties of memory that are not readily detectable through standard behavioral assessments (Dudai et al., 2015).

Taken together, these findings suggest that early post-learning experiences can produce enduring modifications in the state of a memory that influence its future response to reactivation. Because sleep is known to promote memory consolidation through offline reactivation processes, we hypothesized that sleep immediately after learning modifies the future state of a declarative memory, even when no behavioral differences are observed after a prolonged retention interval. We further hypothesized that these sleep-dependent changes remain latent and become evident only when the memory is challenged through a subsequent reactivation procedure.

To test this hypothesis, we conducted two complementary studies. In Study 1, participants learned a declarative memory task and subsequently experienced either a short nap, wakefulness, or an exploratory long-nap condition that included both NREM and REM sleep. Memory was assessed one week later. In Study 2, participants underwent a similar post-learning sleep manipulation, but the memory was reactivated six days after learning using an incomplete reminder previously shown to induce memory updating in human declarative memory under wakefulness conditions (Forcato et al., 2020). Because the primary aim of Study 2 was to determine whether the effects of an initial post-learning nap could be revealed through delayed reactivation, this study focused on the short-nap condition. We reasoned that if sleep modifies the underlying memory, these changes should become evident following reactivation, even when they are not observable during delayed testing alone.

The present study investigated whether post-learning sleep shapes the future fate of a declarative memory. Specifically, we examined whether sleep-dependent consolidation produces enduring modifications that influence how memories respond to subsequent reactivation experiences. By probing the interaction between sleep-dependent consolidation and memory reactivation, we aimed to determine whether sleep leaves latent signatures that remain undetected by conventional memory assessments. Addressing this question may provide new insight into the relationship between sleep-dependent consolidation and memory reconsolidation.

## 2. Materials and Methods

### 2.1 Participants

A total of 107 healthy volunteers (72 women, 35 men; age range: 18-36 years) participated in the study. Participants were undergraduate and graduate students recruited through email advertisements and the laboratory’s social media channels. All participants provided written informed consent prior to participation. The study was approved by the Biomedical Research Ethics Committee of the Instituto Alberto C. Taquini and the Human Ethics Committee of the School of Medicine, University of Buenos Aires, and was conducted in accordance with the Declaration of Helsinki.

Participants were instructed to maintain regular sleep schedules during the week preceding each experimental session, to abstain from alcohol and recreational drugs throughout the study, and to avoid caffeine intake on the day of each experimental session.

Study 1 initially included 62 participants distributed across three experimental conditions: a Short Nap group, a Wake group, and an exploratory Long Nap group designed to obtain a sleep episode containing both NREM and REM sleep. Twenty-one participants were excluded due to incorrect audio recordings (n = 2), inability to fall asleep (n = 7), technical problems (n = 4), failure to understand the task instructions (n = 2), failure to reach REM sleep during the long nap condition (n = 2), or failure to complete all study sessions (n = 4). The final sample consisted of 41 participants (29 women; mean age = 23.9 ± 0.74 years).

Study 2 initially included 45 participants. Seventeen participants were excluded because of medication use during the study (n = 2), inability to fall asleep (n = 6), rehearsal of the learned material outside the laboratory (n = 2), failure to attend scheduled sessions (n = 6), or technical problems (n = 1). The final sample consisted of 28 participants (19 women; mean age = 24.74 ± 1.11 years).

### 2.2 Experimental Design

The study comprised two complementary experiments designed to examine whether post-learning sleep influences the future expression of a declarative memory and its response to later reactivation (Fig 1A).

**Figure 1.**
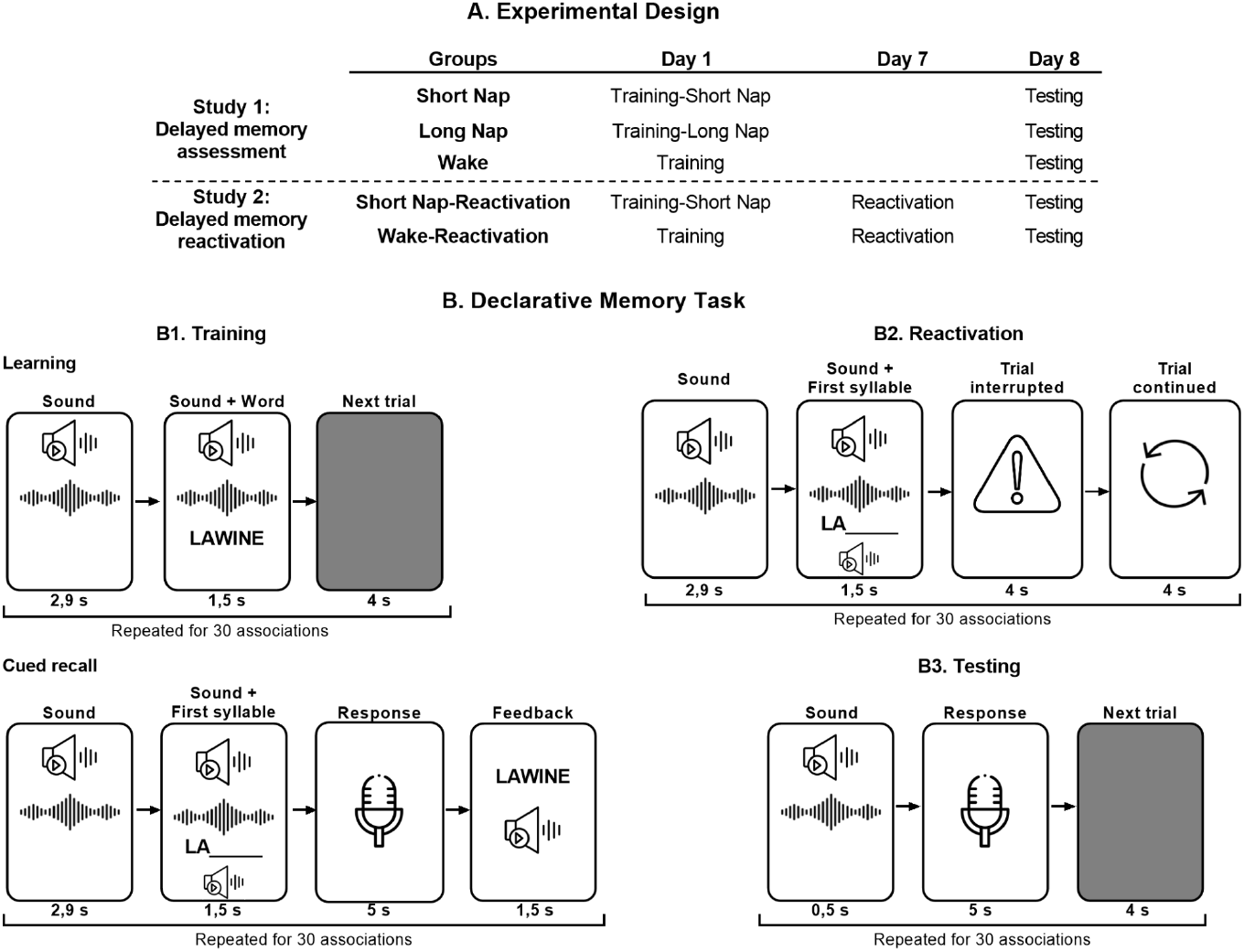
Experimental design and declarative memory task. A) Schematic representation of the experimental procedures used in the two studies. In Study 1, participants were assigned to a Short Nap, Long Nap, or Wake condition after training on Day 1 and completed the memory test on Day 8. In Study 2, participants were assigned to either a Short Nap–Reactivation or Wake–Reactivation condition. Following training on Day 1, all participants underwent a reminder session on Day 7 and were tested on Day 8. B) Declarative memory task. B1. Training: participants learned 30 sound–word associations. During the learning phase, an auditory cue was presented for 2.9 s, followed by the simultaneous presentation of the sound and its corresponding three-syllable word for 1.5 s. During cued recall, the sound and the first syllable of the target word were presented, and participants had 5 s to verbally complete the word, followed by feedback containing the correct response. B2. Reactivation: participants received an incomplete reminder consisting of the original sound cue followed by the first syllable of the associated word. The trial was then interrupted for 4 s and subsequently resumed, inducing prediction error and memory labilization. B3. Testing: during delayed testing, participants heard the original sound cue (0.5 s) and had 5 s to verbally recall the associated word. All phases were repeated for the 30 sound–word associations.

#### Study 1: Delayed Memory Assessment

Participants learned a sound–word association task and were subsequently assigned to one of three post-learning conditions: a short nap (Short Nap group, n = 14), wakefulness (Wake group, n = 14), or an exploratory long-nap condition designed to include both NREM and REM sleep (Long Nap group, n = 13). Memory performance was assessed seven days later.

The Short Nap condition was designed to predominantly contain NREM sleep, whereas the exploratory Long Nap condition provided a longer sleep opportunity that typically included both NREM and REM sleep. The primary aim of Study 1 was to determine whether memory benefits associated with an initial post-learning nap remained detectable after a prolonged retention interval encompassing multiple subsequent sleep–wake cycles. The Long Nap condition was included to explore whether a longer sleep episode containing REM sleep was associated with different patterns of delayed memory performance and sleep–memory relationships.

#### Study 2: Delayed Memory Reactivation

Participants learned the same task and were assigned either to a post-learning short nap (Short Nap-Reactivation group, n = 14) or a wake condition (Wake-Reactivation group, n = 14). Six days after learning, all participants underwent memory reactivation. Memory performance was assessed the following day.

Because the primary objective of Study 2 was to determine whether the effects of an initial post-learning nap could be revealed through delayed reactivation, this experiment focused on the Short Nap condition. We reasoned that if post-learning sleep produces enduring modifications in memory, these changes should become evident following reactivation, even when they are not detectable through delayed memory testing alone.

### 2.3 Experimental Procedure Study 1

The experiment was conducted during daytime hours (11:00-16:00 h) across two sessions separated by one week. Prior to the experimental session, participants assigned to sleep conditions completed an adaptation nap session under identical polysomnographic recording conditions. The adaptation session occurred within the month preceding the experiment and was intended to habituate participants to sleeping in the laboratory environment and with polysomnographic electrodes.

On Day 1, participants completed the Stanford Sleepiness Scale (SSS), learned the declarative memory task, and subsequently underwent one of three post-learning conditions: a short nap, wakefulness, or an exploratory long nap designed to include both NREM and REM sleep. Participants assigned to sleep conditions underwent full polysomnographic recording during the nap period. After the nap, electrodes were removed and participants left the laboratory.

Seven days later (Day 8), participants returned to the laboratory, completed the SSS, and performed the memory test.

#### Study 2

The procedure was similar to that of Study 1 but focused exclusively on the short-nap condition. On Day 1, participants learned the task and subsequently either took a short nap or remained awake.

On Day 7, all participants returned to the laboratory and underwent a memory reactivation procedure. On Day 8, memory performance was assessed.

### 2.4 Declarative Memory Task

The task consisted of learning 30 sound–word associations adapted from Tassone et al. (2020) (Fig 1B). Each environmental sound was semantically associated with a specific Spanish word (e.g., the sound of seeds falling onto the floor paired with the word *PALOMA* [pigeon]). Sounds lasted between 2.85 and 2.94 s (mean duration = 2.9 s) and were presented through headphones. All words consisted of three syllables and were pre-recorded using a female voice.

#### Training

Participants were first exposed to the complete set of 30 sound–word associations. During each trial, a sound was presented for approximately 3 s and then continued while the associated word was simultaneously displayed on the screen and presented auditorily (1.5 s). A black screen followed for 4 s before the next trial (Fig 1.B1).

Learning was subsequently assessed using a cued-recall procedure. Each sound was presented together with the first syllable of the associated word. Participants were instructed to verbally report the complete word when a microphone icon appeared on the screen. After each response period, the correct answer was presented visually and auditorily as feedback.

The training session lasted approximately 15 min. Participants who failed to reach 40% correct responses (12 correct answers) were excluded from further analyses.

#### Memory Reactivation

Memory reactivation was performed only in Study 2 on Day 7.

Participants were informed that the procedure would be identical to the cued-recall task performed during training. Each trial began with the presentation of the sound followed by the first syllable of the associated word. However, contrary to participants’ expectations, the microphone icon never appeared and therefore no response could be given (Fig 1.B2). Instead, the trial was unexpectedly interrupted and a message indicating “Trial interrupted” appeared on the screen, followed by a second message indicating that the experiment was continuing. The procedure was then repeated for the next association until all 30 associations had been presented once.

This reminder procedure was selected because previous studies have shown that incomplete reminders of this type can induce memory updating in human declarative memory under wakefulness conditions (Forcato et al., 2020; Tassone et al., 2020). Consequently, it provided a suitable tool for probing whether post-learning sleep produced latent modifications that influenced the subsequent response of memories to reactivation. Importantly, the reminder differed from the original retrieval condition because participants expected to provide a response that was never allowed, creating a mismatch between expectation and experience.

#### Memory Testing

Memory performance was evaluated using a cued-recall test. Participants heard each sound and were instructed to verbally report the associated word when the microphone icon appeared on the screen (Fig 1.B3). No feedback was provided during testing. Responses were recorded and scored offline.

The primary behavioral outcome was absolute memory change between training and delayed testing. Secondary behavioral measures included the number of correct responses and the distribution of error types at delayed testing. A response was considered correct when the participant accurately recalled the target word associated with a given sound. Synonyms or semantically related words were not accepted as correct responses.

Errors were further classified into three categories. Intralist errors were defined as responses corresponding to another word from the learned list. Confusion errors corresponded to responses that did not belong to the list of learned words. Blank responses were recorded when participants failed to provide any answer.

### 2.5 Polysomnographic Recording and Sleep Analysis

#### 2.5.1 Polysomnography

Sleep was recorded using standard polysomnography, including electroencephalographic (EEG), electrooculographic (EOG), and electromyographic (EMG) recordings acquired with BrainAmp amplifiers (Brain Products GmbH, Munich, Germany). EEG activity was recorded from six scalp electrodes (F3, F4, C3, C4, P3, and P4) placed according to the international 10–20 system. Bilateral mastoid electrodes served as a combined reference. EOG activity was recorded to monitor eye movements and EMG activity was recorded from the chin.

Signals were acquired at a sampling rate of 200 Hz. No online filtering was applied during data acquisition. Offline preprocessing was performed using BrainVision Analyzer (Brain Products GmbH, Germany), and EEG signals were band-pass filtered between 0.16 and 35 Hz.

Polysomnographic recordings were visually scored offline in 30-s epochs according to the criteria of Rechtschaffen and Kales (1968). Sleep stages were classified as wakefulness, stage 1, stage 2, stages 3-4 (slow-wave sleep; SWS), and rapid eye movement (REM) sleep. Conventional sleep parameters were extracted, including total sleep time, sleep onset latency, wake after sleep onset, sleep efficiency, and the amount of time spent in each sleep stage.

#### 2.5.2 Power Spectral Analysis

Power density was calculated separately for all artifact-free epochs of stage 2 sleep, slow-wave sleep (SWS), and REM sleep. Spectral analyses were performed using Fast Fourier Transformations (FFT) with an adapted Hanning window applied to consecutive blocks of 2048 data points (∼8 s) with an overlap of 205 data points (∼0.8 s).

Mean power density values were averaged across electrodes and calculated for the following frequency bands: slow oscillations (0.5-1 Hz), delta (1-4 Hz), theta (4-8 Hz), alpha (8-13 Hz), slow spindle (9-12 Hz) and fast spindle (12-15 Hz). Descriptive power density values for each sleep stage and frequency band are provided in Supporting Information Table S1 (Study 1) and Supporting Information Table S2 (Study 2).

#### 2.5.3 Slow Oscillation and Spindle Detection

Slow oscillations and sleep spindles were detected using SpiSOP (Spindles, Slow Oscillations and Power Spectral Density; RRID:SCR_015673), an open-source toolbox implemented in MATLAB and based on the FieldTrip toolbox (RRID) (Oostenveld et al., 2011; Klinzing et al., 2016; Rudzik et al., 2018; Cha et al., 2020).

For spindle detection, the signal was band-pass filtered around each participant’s individual fast-spindle peak frequency using a 2-Hz bandwidth. The root mean square (RMS) of the filtered signal was calculated and smoothed using a 0.2-s moving average. Spindle events were defined using a threshold of 1.5 standard deviations above the mean RMS signal.

For slow oscillation detection, the signal was filtered between 0.3 and 3.5 Hz. Slow oscillations were identified when the interval between two consecutive positive-to-negative zero crossings ranged between 0.8 and 2 s, corresponding to oscillatory frequencies between 0.5 and 1.25 Hz.

For each participant, the number and density of slow oscillations and sleep spindles were calculated for all electrode locations and averaged across channels. These measures were subsequently used to characterize post-learning sleep and to examine whether individual differences in sleep physiology predicted delayed memory change and the subsequent response of memories to reactivation.

### 2.6 Statistical Analysis

Statistical analyses were performed using SPSS version 21 (IBM Corporation, Armonk, NY, USA). Statistical significance was set at α = 0.05. Unless otherwise specified, all tests were two-tailed.

The primary behavioral outcome was absolute memory change, calculated as the number of correct responses at delayed testing minus the number of correct responses during training. Positive values indicate memory improvement, whereas negative values indicate memory decline across the retention interval.

As secondary behavioral measures, we analyzed the number of correct responses during training and testing, as well as the distribution of error types at delayed testing.

#### Study 1

To determine whether post-learning sleep influenced memory performance after a one-week retention interval, the number of correct responses during training and testing was analyzed using a repeated-measures analysis of variance (ANOVA), with Session (Training, Testing) as a within-subject factor and Group (Short Nap, Long Nap, Wake) as a between-subject factor. Significant interactions were further explored using simple effects analyses.

Absolute memory change was analyzed using a one-way ANOVA with Group (Short Nap, Long Nap, Wake) as a between-subject factor.

#### Study 2

To determine whether post-learning sleep influenced the subsequent response of memories to reactivation, the number of correct responses during training and post-reactivation testing was analyzed using a repeated-measures ANOVA with Session (Training, Testing) as a within-subject factor and Group (Short Nap-Reactivation, Wake-Reactivation) as a between-subject factor. Significant interactions were followed by simple effects analyses.

Absolute memory change was analyzed using independent-samples t-tests comparing the Short Nap-Reactivation and Wake-Reactivation groups.

#### Error Analysis

Errors at testing were classified into three categories. Blank errors were recorded when participants failed to provide any response. Confusion errors corresponded to responses that did not belong to the learned word list. Intralist errors were defined as incorrect responses corresponding to another word from the learned list.

In Study 1, each error type was analyzed using one-way ANOVAs with Group (Short Nap, Long Nap, Wake) as a between-subject factor. In Study 2, error types were analyzed using independent-samples t-tests comparing the Short Nap-Reactivation and Wake-Reactivation groups.

#### Sleep-Memory Associations

To examine whether physiological characteristics of post-learning sleep were associated with memory persistence and the subsequent response of memories to reactivation, Pearson correlation analyses were conducted between sleep parameters and absolute memory change.

For the Short Nap and Short Nap-Reactivation groups, these analyses were performed to evaluate associations between NREM sleep physiology and behavioral outcomes relevant to the primary hypotheses of the study. Additional exploratory analyses were conducted in the Long Nap group to characterize potential associations involving sleep episodes containing both NREM and REM sleep.

Sleep parameters included time spent in each sleep stage, spectral power in the predefined frequency bands, slow oscillation count, and spindle count. Correlation analyses were exploratory in nature and were not corrected for multiple comparisons.

## 3. Results

### 3.1 No detectable behavioral differences after a one-week retention interval

To determine whether post-learning sleep influenced memory performance after a prolonged retention interval, we first analyzed absolute memory change between training (Day 1) and delayed testing (Day 8). Absolute memory change did not differ significantly among the experimental groups (Figure 2.A; one-way ANOVA, F(2,38) = 0.87, p = 0.42). All groups exhibited substantial memory decline over the retention interval (Short Nap: -12.57 ± 1.21; Long Nap: -13.00 ± 0.90; Wake: -11.21 ± 0.69), and memory change differed significantly from zero in each group (all p < 0.001), reflecting the effects of natural forgetting across the week-long delay.

**Figure 2.**
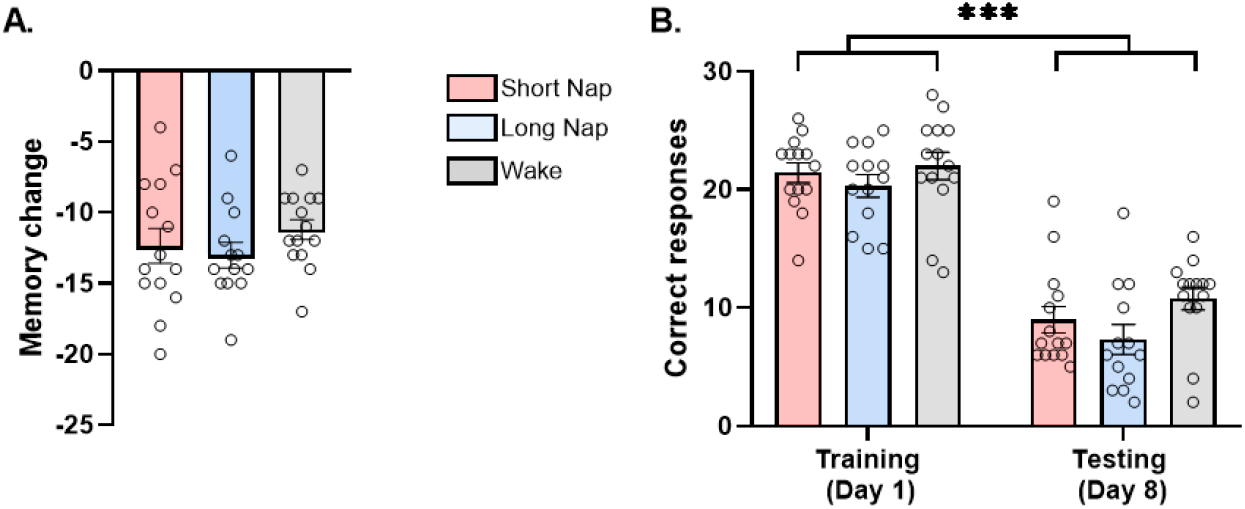
Memory performance in Study 1. A. Memory change (number of correct responses at testing minus the number of correct responses during training) ± SEM. B. Number of correct responses during training on Day 1 ± SEM and number of correct responses during testing on Day 8 ± SEM. * p < 0.05; ** p < 0.01; *** p < 0.001.

Because the Long Nap condition was included for exploratory purposes, we additionally compared the primary experimental conditions

Training performance was comparable across groups (Short Nap: 21.42 ± 0.84; Long Nap: 20.30 ± 0.94; Wake: 22.00 ± 0.55), indicating similar levels of initial learning. A repeated-measures ANOVA revealed a significant main effect of Session (F(1,38) = 485.62, p < 0.001), confirming a marked reduction in memory performance between training and testing. However, neither the main effect of Group (F(2,38) = 1.86, p = 0.17) nor the directly. Consistent with the overall analysis, absolute memory change did not differ between the Short Nap and Wake groups (p = 0.42).

To further characterize memory performance, we analyzed the number of correct responses during training and delayed testing. Group × Session interaction (F(2,38) = 0.94, p = 0.40) reached significance (Figure 2.B).

Finally, error analyses revealed no significant differences among groups in the number of blank, confusion, or intralist errors at delayed testing (all p > 0.10; Table 1).

**Table 1.**
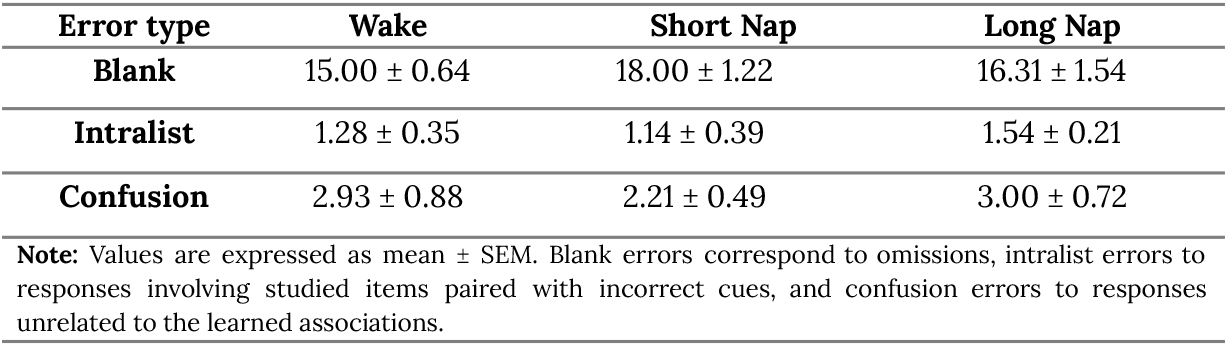
Error types in Study 1.

| Error type | Wake | Short Nap | Long Nap |
| --- | --- | --- | --- |
| Blank | 15.00 ± 0.64 | 18.00 ± 1.22 | 16.31 ± 1.54 |
| Intralist | 1.28 ± 0.35 | 1.14 ± 0.39 | 1.54 ± 0.21 |
| Confusion | 2.93 ± 0.88 | 2.21 ± 0.49 | 3.00 ± 0.72 |

**Note:** Values are expressed as mean ± SEM. Blank errors correspond to omissions, intralist errors to responses involving studied items paired with incorrect cues, and confusion errors to responses unrelated to the learned associations.

Together, these findings indicate that an initial post-learning sleep episode did not produce detectable behavioral advantages after a one-week retention interval, despite substantial forgetting occurring across all groups.

### 3.2 Sleep physiology predicts memory persistence despite the absence of behavioral differences

Although no group differences were observed at delayed testing, several physiological characteristics of post-learning sleep were associated with individual differences in memory retention.

Before examining sleep–memory associations, we verified that the two nap conditions differed in their sleep architecture. As expected from the experimental design, participants in the Long Nap group spent significantly more time in stage 2 sleep, total NREM sleep, and total sleep than participants in the Short Nap group (all p < 0.001; Table 2). No differences were observed in sleep latency, stage 1 sleep, or slow-wave sleep (all p > 0.24).

**Table 2.**
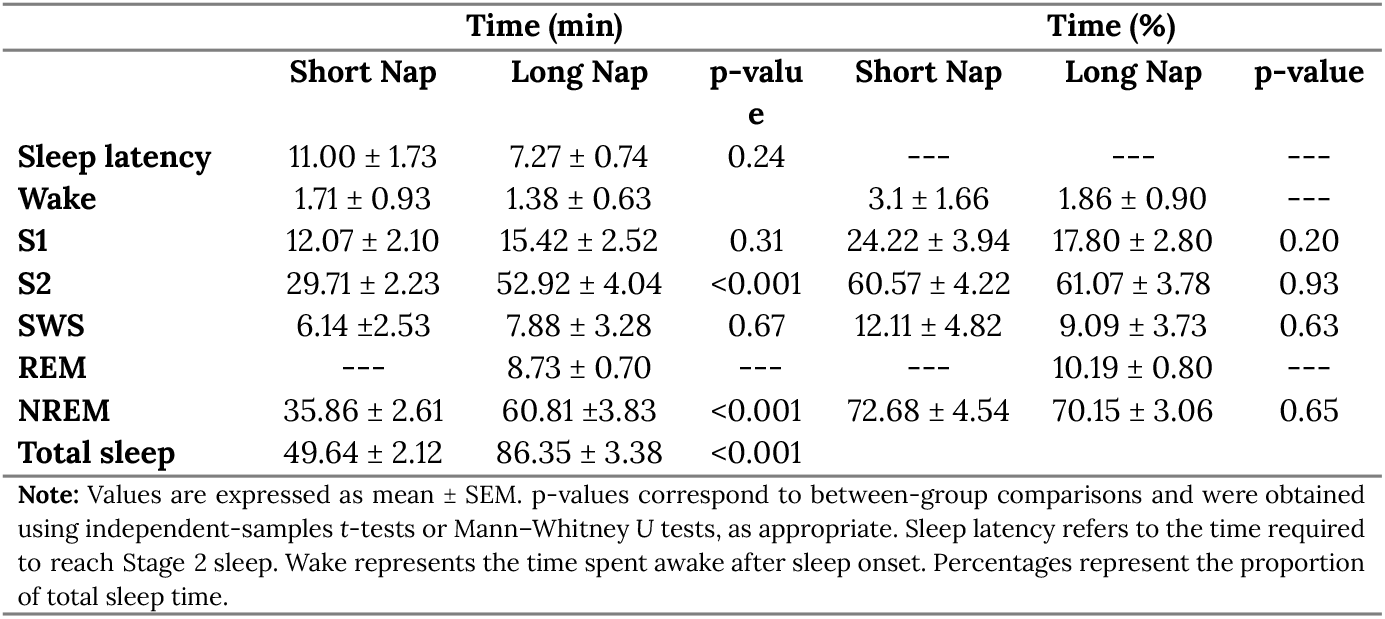
Sleep architecture of the Short Nap and Long Nap groups (Study 1).

|  | Time (min) |  |  | Time (%) |  |  |
| --- | --- | --- | --- | --- | --- | --- |
|  | Short Nap | Long Nap | p-value | Short Nap | Long Nap | p-value |
| **Sleep latency** | $11.00 \pm 1.73$ | $7.27 \pm 0.74$ | 0.24 | --- | --- | --- |
| **Wake** | $1.71 \pm 0.93$ | $1.38 \pm 0.63$ | | $3.1 \pm 1.66$ | $1.86 \pm 0.90$ | --- |
| **S1** | $12.07 \pm 2.10$ | $15.42 \pm 2.52$ | 0.31 | $24.22 \pm 3.94$ | $17.80 \pm 2.80$ | 0.20 |
| **S2** | $29.71 \pm 2.23$ | $52.92 \pm 4.04$ | $<0.001$ | $60.57 \pm 4.22$ | $61.07 \pm 3.78$ | 0.93 |
| **SWS** | $6.14 \pm 2.53$ | $7.88 \pm 3.28$ | 0.67 | $12.11 \pm 4.82$ | $9.09 \pm 3.73$ | 0.63 |
| **REM** | --- | $8.73 \pm 0.70$ | --- | --- | $10.19 \pm 0.80$ | --- |
| **NREM** | $35.86 \pm 2.61$ | $60.81 \pm 3.83$ | $<0.001$ | $72.68 \pm 4.54$ | $70.15 \pm 3.06$ | 0.65 |
| **Total sleep** | $49.64 \pm 2.12$ | $86.35 \pm 3.38$ | $<0.001$ | | | |

**Note:** Values are expressed as mean ± SEM. p-values correspond to between-group comparisons and were obtained using independent-samples t-tests or Mann-Whitney U tests, as appropriate. Sleep latency refers to the time required to reach Stage 2 sleep. Wake represents the time spent awake after sleep onset. Percentages represent the proportion of total sleep time.

#### 3.2.1 NREM sleep physiology and memory persistence

To examine whether NREM sleep physiology was related to memory persistence, correlations were performed between sleep parameters and absolute memory change in the Short Nap group. Longer total sleep time was associated with reduced memory decline across the one-week retention interval (r = 0.47, p = 0.006; Figure 3.A). In addition, greater total sleep time predicted fewer confusion errors at delayed testing (r = -0.42, p = 0.012; Figure 3.B). No significant associations were observed for the remaining sleep stages (all p > 0.06). Power spectral analyses revealed no significant relationships between memory change and slow oscillation power or spindle power across sleep stages (all p > 0.06). Event-based analyses also revealed no significant associations between slow oscillation measures and memory change in the Short Nap group (all p > 0.06). Likewise, no significant associations were observed between memory performance and slow oscillation-spindle coupling measures (all p > 0.27).

**Figure 3.**
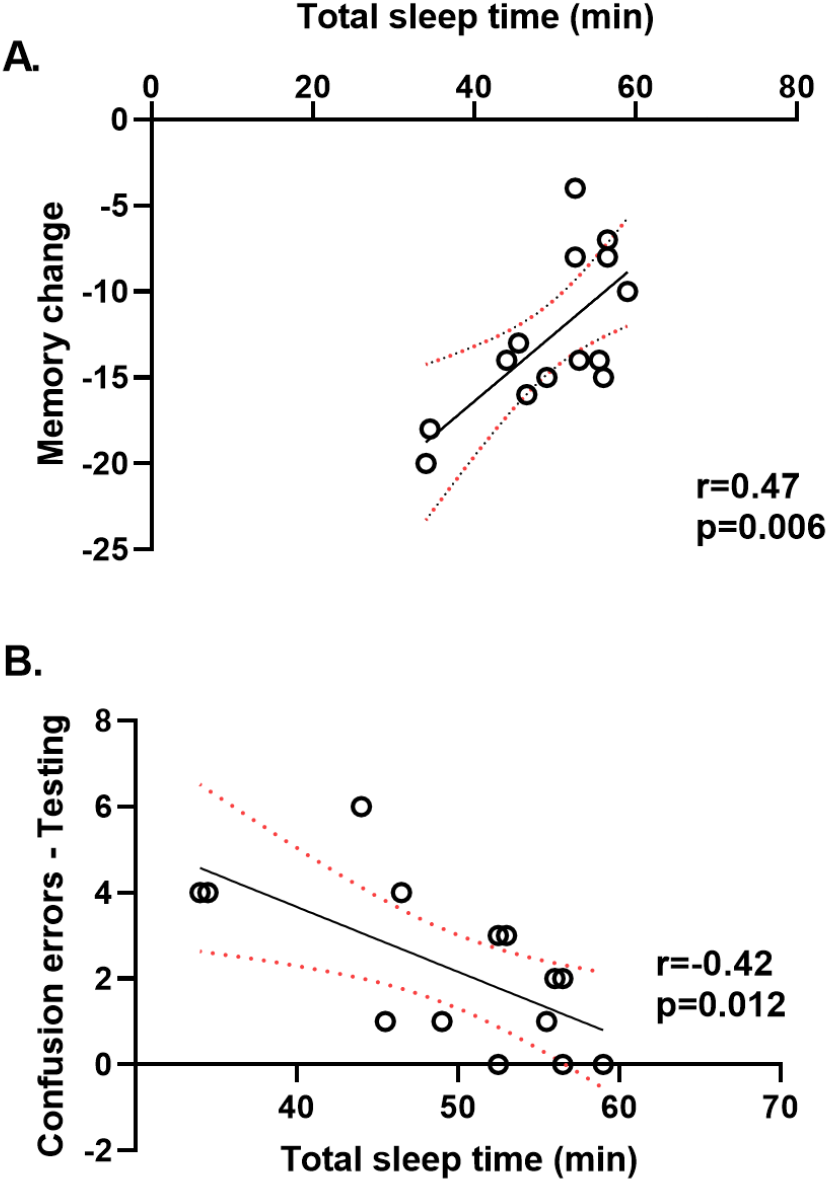
Associations between sleep duration and memory performance in the Short Nap group. A. Correlation between total sleep time and memory change. B. Correlation between total sleep time and the number of confusion errors during memory testing

#### 3.2.2 REM sleep physiology and memory persistence

To further characterize sleep-memory relationships in a longer sleep episode containing both NREM and REM sleep, exploratory correlations were performed between sleep parameters and behavioral outcomes in the Long Nap group. Greater REM sleep duration was associated with increased memory decline (r = -0.34, p = 0.03) and with a higher number of confusion errors at delayed testing (r = 0.34, p = 0.04). No significant relationships were observed between total sleep time and memory performance (all p > 0.23). Power spectral analyses revealed few significant associations with behavioral performance. The only significant association was observed between delta power during SWS and the number of confusion errors, with higher delta power predicting more confusion errors (r = 0.59, p = 0.015). Event-based analyses showed that greater fast spindle density during SWS was associated with reduced memory decline (r = 0.66, p = 0.014), although higher spindle counts were also associated with more confusion errors (r = 0.50, p = 0.048). No significant associations were observed for slow oscillation–spindle coupling measures (all p > 0.27).

Together, these findings indicate that physiological characteristics of post-learning sleep predicted individual differences in memory persistence, even though no behavioral differences were detectable between groups after one week.

### 3.3. Memory reactivation reveals latent effects of post-learning sleep

Because no behavioral differences were observed after one week in Study 1, Study 2 examined whether post-learning sleep produced latent modifications that could be revealed through subsequent memory reactivation.

The primary behavioral outcome, absolute memory change, differed significantly between groups (Figure 4A). Participants in the Short Nap-Reactivation group exhibited a significantly smaller memory decline than participants in the Wake-Reactivation group (-3.78 ± 0.55 vs. -7.57 ± 0.72, respectively; independent-samples t-test, t(26) = 4.20, p < 0.001, Cohen’s d = 1.59), indicating greater memory persistence following post-learning sleep.

**Figure 4.**
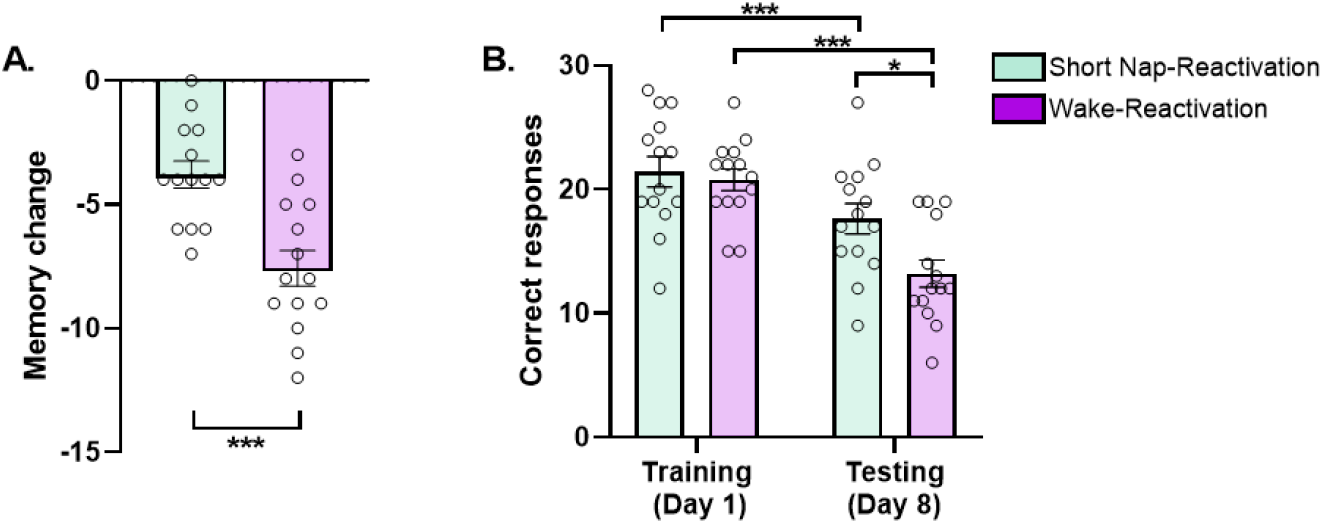
Post-learning sleep reduces memory decline revealed after subsequent reactivation. A. Memory change in the Short Nap-Reactivation and Wake-Reactivation groups. Less negative values indicate reduced memory decline. B. Number of correct responses during training and delayed testing for each group. Error bars represent SEM. Individual data points are shown for each participant. * p < 0.05; ** p < 0.01; *** p < 0.001.

To further characterize this effect, we analyzed the number of correct responses during training and delayed testing using a repeated-measures ANOVA. A significant Group × Session interaction was observed (F(1,26) = 17.65, p < 0.001, η^2^p = 0.40), indicating that the evolution of memory performance from training to testing differed between groups. Simple effects analyses of Group within each Session revealed no significant differences between groups during training (Short Nap-Reactivation: 21.42 ± 1.24; Wake-Reactivation: 20.78 ± 0.88; F(1,26) = 0.18, p = 0.67), confirming comparable levels of initial learning. In contrast, memory performance at delayed testing was significantly higher in the Short Nap-Reactivation group than in the Wake-Reactivation group (17.64 ± 1.22 vs. 13.21 ± 1.09 correct responses, respectively; F(1,26) = 7.29, p = 0.012, η^2^p = 0.22, Figure 4.B).

Simple effects analyses of Session within each Group revealed a significant decline in memory performance between training and testing in both groups (Short Nap-Reactivation: F(1,26) = 35.50, p < 0.001, η^2^p = 0.58; Wake-Reactivation: F(1,26) = 141.20, p < 0.001, η^2^p = 0.84). However, the magnitude of forgetting was substantially smaller in participants who slept after learning. Finally, error analyses revealed no significant differences between groups in the number of blank, intralist, or confusion errors at delayed testing (Table 3; all p > 0.051).

**Table 3.** Error types in Study 2.

| Type of error | Short Nap Reactivation | Wake Reactivation |
| --- | --- | --- |
| Blank | $8.57 \pm 1.16$ | $12.43 \pm 1.49$ |
| Intralist | $1.17 \pm 0.27$ | $1.36 \pm 0.29$ |
| Confusion | $2.00 \pm 0.69$ | $3.00 \pm 0.72$ |

**Note:** Values are expressed as mean ± SEM. Blank errors correspond to omissions, intralist errors to responses involving studied items paired with incorrect cues, and confusion errors to responses unrelated to the learned associations.

Together, these findings indicate that post-learning sleep produced enduring modifications in the memory trace that were not detectable through delayed testing alone, but became behaviorally evident following subsequent memory reactivation. These results suggest that sleep influences not only the persistence of declarative memories, but also their subsequent responsiveness to reactivation experiences.

### 3.4. NREM sleep physiology predicts the response of memories to subsequent reactivation

To identify physiological characteristics of post-learning sleep associated with the subsequent response of memories to reactivation, we examined correlations between sleep parameters and absolute memory change in the Short Nap-Reactivation group. Descriptive sleep characteristics of the Short Nap-Reactivation group are presented in Table 4.

**Table 4.** Sleep architecture of the Short Nap-Reactivation group.

|  | Short Nap-Reactivation |  |
| --- | --- | --- |
|  | Time (min) | Time (%) |
| **Sleep latency** | 8.36 ± 1.95 | — |
| **Wake** | 2.39 ± 0.66 | 5.03 ± 1.40 |
| **S1** | 10.36 ± 1.95 | 21.62 ± 3.12 |
| **S2** | 26.82 ± 1.77 | 58.88 ± 3.72 |
| **SWS** | 6.64 ± 1.96 | 14.46 ± 4.10 |
| **NREM** | 33.46 ± 2.15 | 73.34 ± 4.09 |
| **Total sleep** | 46.21 ± 2.50 | --- |

**Note.** Values are expressed as mean ± SEM. Sleep latency refers to the time required to reach Stage 2 sleep. Wakefulness refers to the time spent awake after sleep onset. Percentages are expressed relative to total sleep time.

Among conventional sleep measures, greater time spent in SWS (SWS) was associated with reduced memory decline (all p < 0.05; Figure 5A, B). No significant associations were observed between memory change and time spent in other sleep stages (all p > 0.061). Spectral analyses revealed that higher slow oscillation power density (0.5–1 Hz) during NREM sleep was associated with reduced memory decline (r = 0.31, p = 0.038; Figure 5C). No significant correlations were observed for other frequency bands or sleep stages (all p > 0.06). Event-based analyses showed that reduced memory decline was associated with both a greater number of slow oscillations and higher slow oscillation density during NREM sleep (r = 0.34, p = 0.026 and r = 0.53, p = 0.003, respectively; Figure 5D, E). Similarly, a greater number of fast spindles during SWS was associated with reduced memory decline (r = 0.56, p = 0.021; Figure 5F). No significant associations were observed between memory performance and slow oscillation–spindle coupling measures (all p > 0.37).

**Figure 5.**
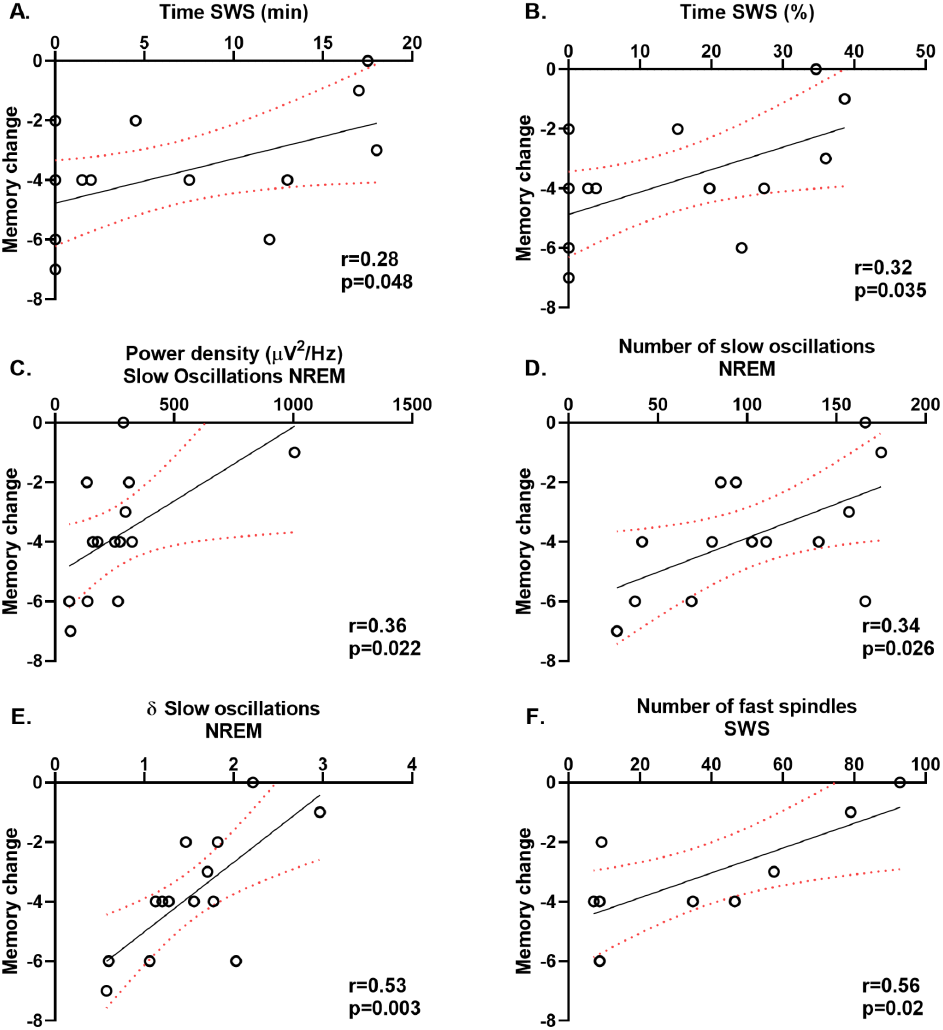
Slow-wave sleep and NREM oscillatory activity are associated with reduced memory decline following reactivation. A. Correlation between SWS duration (min) and memory change. B. Correlation between SWS duration (%) and memory change. C. Correlation between NREM slow oscillation power density (0.5–1 Hz) and memory change. D. Correlation between the number of slow oscillations during NREM sleep and memory change. E. Correlation between slow oscillation density during NREM sleep and absolute memory change. F. Correlation between the number of fast spindles during SWS and memory change. Each dot represents one participant from the Short Nap-Reactivation group.

Together, these findings indicate that individual differences in NREM sleep physiology, particularly SWS duration, slow oscillatory activity, and spindle expression, predict the subsequent response of memories to reactivation.

## Discussion

Although an initial post-learning nap did not produce detectable behavioral advantages after a one-week retention interval, these latent sleep-dependent modifications became behaviorally evident when memories were subsequently reactivated. Although an initial post-learning nap did not produce detectable behavioral advantages after a one-week retention interval, these latent effects became evident when memories were subsequently reactivated. Moreover, physiological characteristics of NREM sleep predicted the magnitude of this post-reactivation memory benefit. Together, these findings suggest that sleep-dependent consolidation influences the future responsiveness of memories to reactivation experiences.

The first important observation was that no behavioral differences were detectable one week after learning. At first glance, this result could be interpreted as evidence that post-learning sleep did not exert lasting effects on memory. However, such an interpretation would be inconsistent with both the physiological findings of Study 1 and the reactivation findings of Study 2. Instead, our results suggest that sleep-dependent changes may persist in latent forms that are not readily captured by conventional delayed memory assessments. Indeed, all groups exhibited substantial forgetting across the retention interval, suggesting that behavioral differences initially produced during consolidation may progressively diminish over time as memories continue to evolve across subsequent cycles of wakefulness and sleep.

Importantly, the idea that the behavioral expression of memory does not necessarily reveal its underlying state is consistent with previous findings from the reconsolidation literature. For example, Forcato et al. (2013) demonstrated that repeated reactivation procedures strengthened relatively recent declarative memories but failed to produce comparable effects when memories were reactivated seven days after learning. These findings suggested that memory age acts as a limiting condition for memory strengthening through reconsolidation. In the present study, however, delayed reactivation revealed differences between memories encoded before sleep and memories encoded before wakefulness that were no longer detectable through delayed testing alone. Thus, although behavioral differences were no longer detectable after one week, the subsequent response to reactivation revealed that the memories were not in the same state.

A similar conclusion emerges from the work of Fernández et al. (2016), who showed that a stressful experience administered shortly after learning altered the subsequent response of a seven-day-old declarative memory to reactivation. In that study, memory strengthening effects that were absent in control participants emerged when memories had previously been exposed to post-learning stress. Together, these findings suggest that experiences occurring during early stages of consolidation can shape the future fate of a memory trace, influencing how it responds to later reactivation experiences. The present results extend this principle to sleep, indicating that post-learning sleep can produce long-lasting modifications that remain behaviorally silent until the memory is subsequently challenged. Notably, both studies suggest that experiences occurring shortly after learning may alter the future trajectory of memory processing without necessarily producing immediately detectable behavioral consequences.

Sleep represents a particularly relevant candidate for producing such enduring modifications because it promotes offline memory processing through spontaneous reactivation mechanisms. According to the Active Systems Consolidation framework, recently encoded memories are repeatedly reactivated during NREM sleep through coordinated interactions between cortical slow oscillations, thalamocortical sleep spindles, and hippocampal sharp-wave ripples (Diekelmann & Born, 2010; Rasch & Born, 2013; Klinzing et al., 2019). Traditionally, these processes have been interpreted as strengthening memory representations and improving later recall. The present findings suggest a broader interpretation. Rather than simply increasing memory strength, post-learning sleep may alter the state of the memory itself, producing enduring changes that remain behaviorally silent yet influence how memories respond to future reactivation experiences.

This interpretation aligns closely with theoretical proposals suggesting that sleep reorganizes memory representations rather than merely stabilizing them (Lewis & Durrant, 2011; Stickgold & Walker, 2013). Through repeated offline reactivation, memories may become progressively integrated within existing knowledge networks, acquiring properties that influence their future accessibility, flexibility, and susceptibility to modification. Under this view, sleep-dependent consolidation does not simply determine how much is remembered. Instead, it influences what memories become.

The physiological findings support this interpretation. In Study 2, greater time spent in SWS, higher slow oscillation power during NREM sleep, increased slow oscillation density, and a greater number of fast sleep spindles were all associated with reduced memory decline following delayed reactivation. These findings are highly consistent with the proposed role of slow oscillations and sleep spindles in supporting hippocampal-neocortical communication and long-term declarative memory consolidation (Mölle et al., 2009; Staresina et al., 2015; Helfrich et al., 2018). Importantly, however, these physiological markers predicted not immediate memory performance, but rather the subsequent responsiveness of memories to reactivation. This observation extends traditional views of sleep-dependent consolidation by suggesting that NREM sleep physiology contributes not only to memory stabilization but also to the future capacity of memories to benefit from later reactivation experiences.

Interestingly, Study 1 revealed several physiological associations despite the absence of behavioral group differences. In particular, longer total sleep time was associated with reduced forgetting and fewer confusion errors one week later. These findings provide additional support for the idea that sleep-related processes continue to influence memory persistence even when their effects are no longer readily detectable at the behavioral level. Nevertheless, the strongest physiological associations emerged in Study 2, when memory was challenged through reactivation, suggesting that delayed reactivation may provide a more sensitive assay of sleep-dependent memory modifications than delayed testing alone.

The exploratory long-nap condition provided preliminary observations regarding the potential contribution of REM sleep to long-term memory outcomes. Although no behavioral differences emerged at the group level, greater REM sleep duration was associated with increased memory decline and a higher number of confusion errors one week later. These findings may initially appear counterintuitive but are consistent with proposals suggesting that REM sleep contributes to memory integration, abstraction, and generalization rather than to the preservation of precise episodic details (Sterpenich et al., 2014; Payne, 2014; Lewis et al., 2018). Under this interpretation, increased confusion errors may not reflect simple forgetting but rather the emergence of more generalized memory representations that preserve semantic relationships at the expense of exact item-specific information.

This interpretation is further supported by studies suggesting that REM sleep may promote the integration, abstraction, and generalization of newly acquired information, whereas NREM sleep may preferentially support the stabilization of specific episodic traces (Lewis et al., 2018; Payne, 2014; Sterpenich et al., 2014). In line with this view, Kaida et al. (2023) reported that REM-rich sleep was associated with increased memory distortions, whereas NREM-rich sleep preferentially supported memory stabilization. Together, these observations are compatible with the notion that NREM and REM sleep contribute complementary processes during memory formation, with NREM sleep promoting the stabilization and persistence of specific memory traces and REM sleep facilitating their integration within broader associative networks. However, because the long-nap condition was included for exploratory purposes and was not incorporated into Study 2, the present data do not allow direct conclusions regarding the contribution of REM sleep to the reactivation effects observed here.

Several limitations should be acknowledged. First, the correlational nature of the physiological analyses precludes causal conclusions regarding the role of specific sleep oscillations in producing the observed effects. Second, the exploratory long-nap condition was not included in the reactivation study, preventing direct comparisons between NREM- and REM-rich sleep episodes with respect to their influence on later memory reactivation. Consequently, it remains unclear whether REM-rich sleep would produce similar, weaker, or qualitatively different effects on subsequent memory reactivation. Future studies combining targeted manipulations of sleep physiology with delayed memory reactivation procedures will be necessary to clarify the mechanisms underlying these effects.

In conclusion, the present findings indicate that post-learning sleep does not simply determine whether memories are retained over time. Rather, sleep induces enduring modifications that shape the future expression of memory and its responsiveness to later reactivation experiences. These effects remain largely invisible to conventional delayed memory tests but become evident when memories are challenged through reactivation. Together, these findings provide evidence that sleep-dependent consolidation and memory reconsolidation are intimately linked processes and suggest that sleep influences not only how much we remember, but also what our memories ultimately become.

## Author contribution

Malen D. Moyano: Conceptualization; methodology; investigation; visualization; writing original draft; writing - review and editing; formal analysis; data curation. Micaela S. Lombardi: Investigation; writing - review and editing. Aylin A. Vázquez Chenlo: Software; writing - review and editing. Luis I. Brusco: Resources, writing - review and editing. Cecilia Forcato: Conceptualization, methodology; writing original draft; writing - review and editing; resources; project administration; supervision.

## Data availability statement

All behavioral and electrophysiological data generated and analyzed during this study will be made publicly available in an open-access repository upon acceptance of the manuscript. Repository details and persistent identifiers (DOI) will be provided before publication to ensure unrestricted access to the data supporting the findings of this study.

## Use of Artificial Intelligence

The authors used ChatGPT (OpenAI) exclusively to assist with translation and English language editing of the manuscript, as the authors’ native language is Spanish. ChatGPT was also used to obtain general suggestions for improving the clarity and visual organization of the experimental protocol figure. No AI tools were used for data analysis, interpretation of results, or generation of scientific conclusions. The authors reviewed and edited all AI-assisted text and figure-related suggestions and take full responsibility for the final content of the manuscript.

## Supporting Information

**Table S1.** Power density (µV^2^/Hz) in the frequency bands of interest during each sleep stage, expressed as mean ± SEM for Short Nap and Long Nap groups.

|  | Short Nap | Long Nap |
| --- | --- | --- |
| **NREM** |  |  |
| Slow oscillations (0.5 - 1 Hz) | 239.68 ± 60.99 | 249.99 ± 47.96 |
| Delta (1 - 4 Hz) | 57.37 ± 11.70 | 61.24 ± 12.02 |
| Theta (4 -8 Hz) | 7.37 ± 0.74 | 7.96 ± 0.76 |
| Slow spindles (9-12Hz) | 3.14 ± 0.39 | 2.93 ± 0.34 |
| Fast spindles (12-15Hz) | 2.68 ± 0.49 | 2.42 ± 0.36 |
| **SWS** |  |  |
| Slow oscillations (0.5 - 1 Hz) | 720.50 ± 123.26 | 787.48 ± 86.55 |
| Delta (1 - 4 Hz) | 146.54 ± 24.57 | 154.88 ± 19.61 |
| Theta (4 -8 Hz) | 9.92 ± 1.11 | 10.75 ± 0.86 |
| Slow spindles (9-12Hz) | 3.17 ± 0.52 | 2.60 ± 0.28 |
| Fast spindles (12-15Hz) | 2.07 ± 0.73 | 1.58 ± 0.17 |
| **S2** |  |  |
| Slow oscillations (0.5 - 1 Hz) | 143.04 ± 19.99 | 166.20 ± 19.58 |
| Delta (1 - 4 Hz) | 40.31 ± 4.71 | 44.53 ± 4.62 |
| Theta (4 -8 Hz) | 7.02 ± 0.70 | 7.78 ± 0.72 |
| Slow spindles (9-12Hz) | 3.19 ± 0.40 | 2.97 ± 0.35 |
| Fast spindles (12-15Hz) | 2.78 ± 0.49 | 2.61 ± 0.43 |
| **REM** |  |  |
| Theta (4-8Hz) |  | 3.66 ± 0.43 |

**Table S2.**
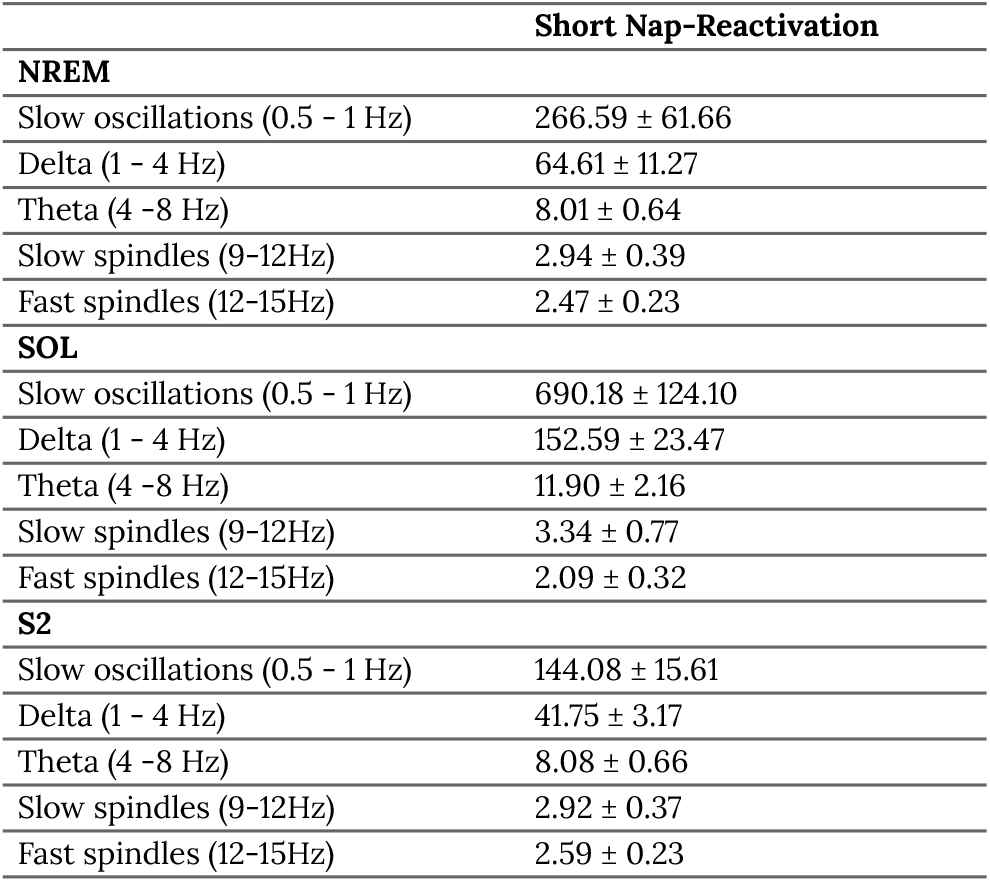
Power density (µV^2^/Hz) in the frequency bands of interest during each sleep stage, expressed as mean ± SEM for Short Nap-Reactivation.

## References

Cha, K. S., Kim, T. J., Jun, J. S., Byun, J. I., Sunwoo, J. S., Shin, J. W., Kim, K. H., Lee, S. K., & Jung, K. Y. (2020). Impaired slow oscillation, sleep spindle, and slow oscillation-spindle coordination in patients with idiopathic restless legs syndrome. Sleep medicine, 66, 139–147. 10.1016/j.sleep.2019.09.021

Diekelmann, S., & Born, J. (2010). The memory function of sleep. Nature Reviews. Neuroscience, 11(2), 114–126. 10.1038/nrn2762

Dudai Y. (2004). The neurobiology of consolidations, or, how stable is the engram?. Annual review of psychology, 55, 51–86. 10.1146/annurev.psych.55.090902.142050

Dudai Y. (2006). Reconsolidation: the advantage of being refocused. Current opinion in neurobiology, 16(2), 174–178. 10.1016/j.conb.2006.03.010

Dudai, Y., Karni, A., & Born, J. (2015). The Consolidation and Transformation of Memory. Neuron, 88(1), 20–32. 10.1016/j.neuron.2015.09.004

Ellenbogen, J. M., Payne, J. D., & Stickgold, R. (2006). The role of sleep in declarative memory consolidation: passive, permissive, active or none?. Current opinion in neurobiology, 16(6), 716–722. 10.1016/j.conb.2006.10.006

Fernández, R. S., Bavassi, L., Forcato, C., & Pedreira, M. E. (2016). The dynamic nature of the reconsolidation process and its boundary conditions: Evidence based on human tests. Neurobiology of learning and memory, 130, 202–212. 10.1016/j.nlm.2016.03.001

Forcato, C., Fernandez, R. S., & Pedreira, M. E. (2013). The role and dynamic of strengthening in the reconsolidation process in a human declarative memory: what decides the fate of recent and older memories?. PloS one, 8(4), e61688. 10.1371/journal.pone.0061688

Forcato, C., Klinzing, J. G., Carbone, J., Radloff, M., Weber, F. D., Born, J., & Diekelmann, S. (2020). Reactivation during sleep with incomplete reminder cues rather than complete ones stabilizes long-term memory in humans. Communications biology, 3(1), 733. 10.1038/s42003-020-01457-4

Forcato, C., Rodríguez, M. L., & Pedreira, M. E. (2011). Repeated labilization-reconsolidation processes strengthen declarative memory in humans. PloS one, 6(8), e23305. 10.1371/journal.pone.0023305

Frankland, P. W., & Bontempi, B. (2005). The organization of recent and remote memories. Nature reviews. Neuroscience, 6(2), 119–130. 10.1038/nrn1607

Gais, S., Albouy, G., Boly, M., Dang-Vu, T. T., Darsaud, A., Desseilles, M., Rauchs, G., Schabus, M., Sterpenich, V., Vandewalle, G., Maquet, P., & Peigneux, P. (2007). Sleep transforms the cerebral trace of declarative memories. Proceedings of the National Academy of Sciences of the United States of America, 104(47), 18778–18783.10.1073/pnas.0705454104

Helfrich, R. F., Mander, B. A., Jagust, W. J., Knight, R. T., & Walker, M. P. (2018). Old Brains Come Uncoupled in Sleep: Slow Wave-Spindle Synchrony, Brain Atrophy, and Forgetting. Neuron, 97(1), 221–230.e4. 10.1016/j.neuron.2017.11.020

Kaida, K., Mori, I., Kihara, K., & Kaida, N. (2023). The function of REM and NREM sleep on memory distortion and consolidation. Neurobiology of learning and memory, 204, 107811. 10.1016/j.nlm.2023.107811

Klinzing, J. G., Mölle, M., Weber, F., Supp, G., Hipp, J. F., Engel, A. K., & Born, J. (2016). Spindle activity phase-locked to sleep slow oscillations. NeuroImage, 134, 607–616. 10.1016/j.neuroimage.2016.04.031

Klinzing, J. G., Niethard, N., & Born, J. (2019). Mechanisms of systems memory consolidation during sleep. Nature Neuroscience, 22(10), 1598–1610. 10.1038/s41593-019-0467-3

Latchoumane, C. V., Ngo, H. V., Born, J., & Shin, H. S. (2017). Thalamic Spindles Promote Memory Formation during Sleep through Triple Phase-Locking of Cortical, Thalamic, and Hippocampal Rhythms. Neuron, 95(2), 424–435.e6. 10.1016/j.neuron.2017.06.025

Lewis, P. A., & Durrant, S. J. (2011). Overlapping memory replay during sleep builds cognitive schemata. Trends in cognitive sciences, 15(8), 343–351. 10.1016/j.tics.2011.06.004

Lewis, P. A., Knoblich, G., & Poe, G. (2018). How Memory Replay in Sleep Boosts Creative Problem-Solving. Trends in cognitive sciences, 22(6), 491–503. 10.1016/j.tics.2018.03.009

Mölle, M., Eschenko, O., Gais, S., Sara, S. J., & Born, J. (2009). The influence of learning on sleep slow oscillations and associated spindles and ripples in humans and rats. The European journal of neuroscience, 29(5), 1071–1081. 10.1111/j.1460-9568.2009.06654.x

Nader, K., & Hardt, O. (2009). A single standard for memory: the case for reconsolidation. Nature reviews. Neuroscience, 10(3), 224–234. 10.1038/nrn2590

Nader, K., Schafe, G. E., & Le Doux, J. E. (2000). Fear memories require protein synthesis in the amygdala for reconsolidation after retrieval. Nature, 406(6797), 722–726. 10.1038/35021052

Oostenveld, R., Fries, P., Maris, E., & Schoffelen, J. M. (2011). FieldTrip: Open Source Software for Advanced Analysis of MEG, EEG, and Invasive Electrophysiological Data. Comput Intell Neurosci. 156869. 10.1155/2011/156869

Payne J. D. (2014). Seeing the forest through the trees. Sleep, 37(6), 1029–1030. 10.5665/sleep.3750

Rasch, B., & Born, J. (2013). About sleep’s role in memory. Physiological reviews, 93(2), 681–766. 10.1152/physrev.00032.2012

Rechtschaffen, A., & Kales, A. (1968). A Manual of Standardized Terminology, Technique and Scoring System for Sleep Stages of Human Sleep. Brain Information Service, Brain Information Institute, UCLA: Los Angeles, CA, USA.

Rudzik, F., Thiesse, L., Pieren, R., Wunderli, J. M., Brink, M., Foraster, M., Héritier, H., Eze, I. C., Garbazza, C., Vienneau, D., Probst-Hensch, N., Röösli, M., & Cajochen, C. (2018). Sleep spindle characteristics and arousability from nighttime transportation noise exposure in healthy young and older individuals. Sleep, 41(7), 10.1093/sleep/zsy077. https://doi.org/10.1093/sleep/zsy077

Squire, L. R., & Alvarez, P. (1995). Retrograde amnesia and memory consolidation: a neurobiological perspective. Current opinion in neurobiology, 5(2), 169–177. 10.1016/0959-4388(95)80023-9

Staresina, B. P., Bergmann, T. O., Bonnefond, M., van der Meij, R., Jensen, O., Deuker, L., Elger, C. E., Axmacher, N., & Fell, J. (2015). Hierarchical nesting of slow oscillations, spindles and ripples in the human hippocampus during sleep. Nature neuroscience, 18(11), 1679–1686. 10.1038/nn.4119

Sterpenich, V., Schmidt, C., Albouy, G., Matarazzo, L., Vanhaudenhuyse, A., Boveroux, P., Degueldre, C., Leclercq, Y., Balteau, E., Collette, F., Luxen, A., Phillips, C., & Maquet, P. (2014). Memory reactivation during rapid eye movement sleep promotes its generalization and integration in cortical stores. Sleep, 37(6), 1061–1075B. 10.5665/sleep.3762

Stickgold, R., & Walker, M. P. (2013). Sleep-dependent memory triage: evolving generalization through selective processing. Nature neuroscience, 16(2), 139–145. 10.1038/nn.3303

Suzuki, A., Josselyn, S. A., Frankland, P. W., Masushige, S., Silva, A. J., & Kida, S. (2004). Memory reconsolidation and extinction have distinct temporal and biochemical signatures. The Journal of neuroscience : the official journal of the Society for Neuroscience, 24(20), 4787–4795. 10.1523/JNEUROSCI.5491-03.2004

Tassone, L. M., Urreta Benítez, F. A., Rochon, D., Martínez, P. B., Bonilla, M., Leon, C. S., Muchnik, C., Solis, P., Medel, N., Kochen, S., Brusco, L. I., Moyano, M. D., & Forcato, C. (2020). Memory reconsolidation as a tool to endure encoding deficits in elderly. PloS one, 15(8), e0237361. 10.1371/journal.pone.0237361

Wang, S. H., de Oliveira Alvares, L., & Nader, K. (2009). Cellular and systems mechanisms of memory strength as a constraint on auditory fear reconsolidation. Nature neuroscience, 12(7), 905–912. 10.1038/nn.2350

